# Transcription Factor NRF2 is Activated by Erythrophagocytosis of Oxidized Red Blood Cell Products and Suppresses the IL-12-IFNg-IL-10 Axis in a Murine Model of Hyperinflammatory Disease

**DOI:** 10.1101/2023.12.12.571271

**Authors:** Paul M. Gallo, Em Elliott, Grace Ford, Chhanda Biswas, Jihwan Kim, Jadyn Wheaton, Connie Jiang, Niansheng Chu, Portia A. Kreiger, Michele Lambert, Edward M. Behrens

## Abstract

Hyperinflammatory diseases including macrophage activation syndrome (MAS) and hemophagocytic lymphohistocytosis (HLH) are characterized by multi-lineage cytopenias, hypercytokinemia, and tissue hemophagocytosis. However, the mechanisms by which erythrophagocytosis mediates iron metabolism and regulates the hyperactive immune response remain unclear. The transcription factor NRF2 is an important sensor for inflammatory and redox distress. The targets of NRF2 are antioxidant response elements responsible for transcription of genes related to restoration of redox homeostasis within the cell. Here we demonstrate that mice with CpG-induced MAS have evidence of systemic oxidative and nitrosative distress – including increased serum nitric oxide and elevated systemic lipid peroxidation. In this model, NRF2 knockout mice develop significantly worse organomegaly, hypercytokinemia, and reticulocytosis. NRF2 knockout mice have unexpected exacerbation in the cytokines that are central to hyperinflammatory physiology – namely IL-12, IFN gamma (IFNg), and IL-10. *In vitro* we demonstrate that oxidized red blood cell products and heme itself suppress IL-12 protein production and transcription from bone marrow derived dendritic cells in a NRF2-dependent manner. Together our studies demonstrate that erythrophagocytosis of oxidized red blood cell products suppresses the Il-12-IFNg-IL-10 axis which drives hyperinflammation in murine hyperinflammation.

**Graphical abstract:** 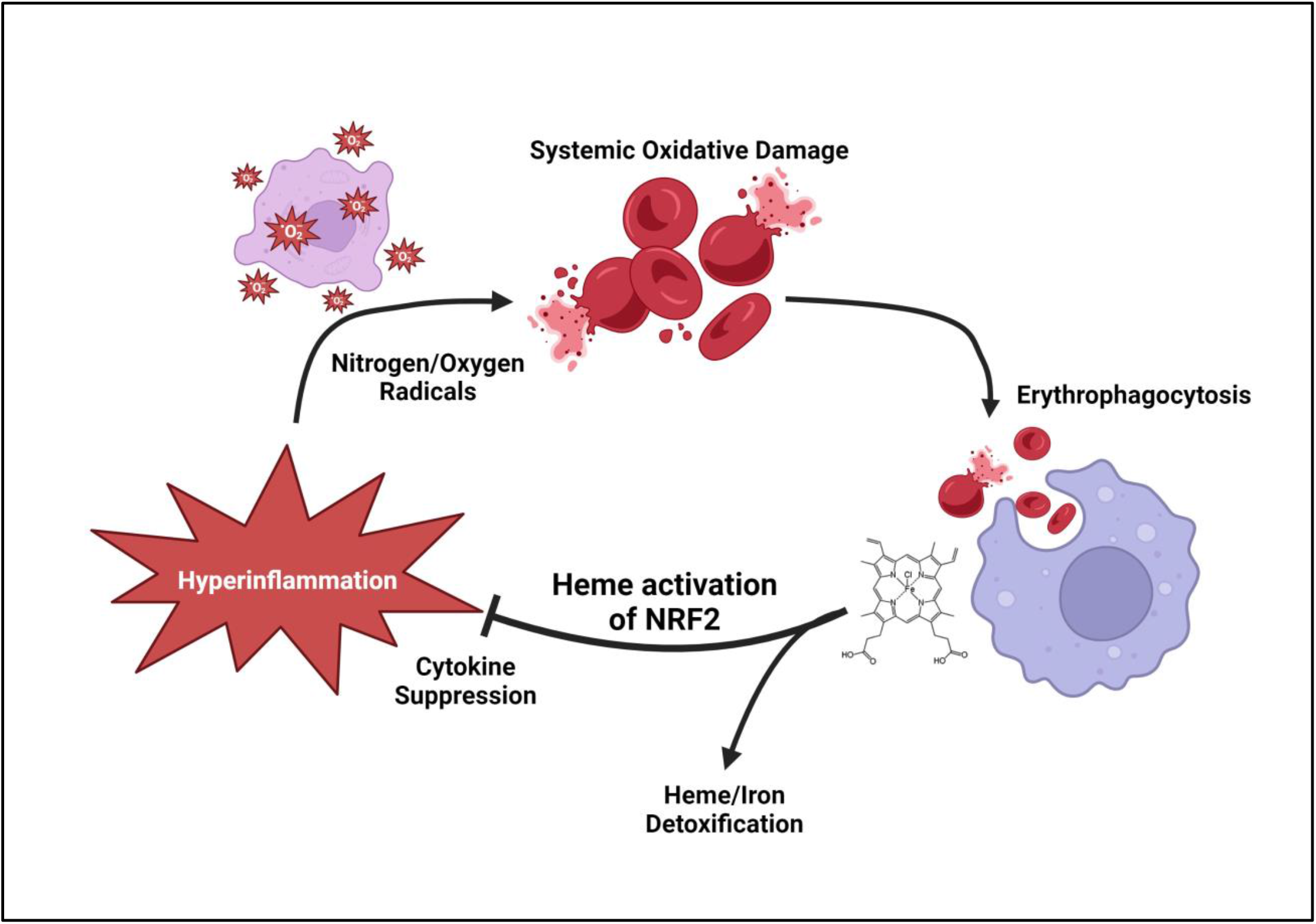

Created with BioRender

## 1. Introduction

Macrophage activation syndrome (MAS) and hemophagocytic lymphohistocytosis (HLH) are cytokine storm syndromes that represent a spectrum of hyperinflammatory physiology. These conditions are clinically characterized by high fevers, multi-lineage cytopenias, hypercytokinemia, and multisystem organ failure [1]. On closer inspection of the classic clinical criteria for HLH, a common theme emerges of iron metabolism and red blood cell redox biology. Anemia is common in MAS/HLH, high serum ferritin is sensitive for HLH physiology, tissue hemophagocytosis is a unique marker of hyperinflammatory disease, and splenomegaly (largely due to expansion of iron recycling red pulp macrophages) is nearly universal in MAS/HLH. Beyond the classic criteria of MAS/HLH, observations of high heme oxygenase 1 (HO1) expression [2] and elevated soluble CD163 (haptoglobin receptor) [3] also point towards maladaptive iron metabolism in systemic hyperinflammation.

Findings from our lab has demonstrated that HO1 is not only an interesting biomarker for hyperinflammatory processes, but also an important mediator of disease. For example, we have found that the small molecule monomethyl fumarate (MMF) is able to ameliorate CpG-induced murine hyperinflammation in a HO1-dependent and HO1-independent manner [4]. MMF and its relative dimethyl fumarate are potent inducers of transcription factor NRF2, which is well established as a critical sensor and regulator of antioxidant cellular pathways [5-7]. Experiments from others have shown that RBC products activate NRF2, resulting in relative immunosuppression [8, 9]. We therefore hypothesized that murine hyperinflammation would be exacerbated by NRF2 deficiency and that hemophagocytosis of RBC products acts to regulate proinflammatory cytokine production in a NRF2-dependent manner.

Here we find that NRF2 deficiency intensifies CpG-induced murine hyperinflammation, leading to worsened systemic oxidative/nitrosative distress and exacerbation of the IL-12-IFNg-IL10 hypercytokine axis. We demonstrate that products of oxidized RBCs are present *in vivo* after induction of hyperinflammation and that these products suppress hyperinflammatory IL-12 cytokine production through a NRF2-dependent mechanism *in vitro*.

## 2. Methods

### 2.1 Mice

This study was carried out in the animal facility of Children’s Hospital of Philadelphia (Philadelphia, Pennsylvania) and was approved by the Institutional Animal Care and Use Committee (protocol no. 921). C57BL/6 (wild-type [WT]) and NRF2 KO mice were purchased from The Jackson Laboratory and bred in our facility. The HMOX1fl/fl mice were a generous gift from Dr. Zoltan Arany (Perelman School of Medicine, University of Pennsylvania). HMOX1fl/fl mice were crossbred with LysMcre mice in our facility and genotyped regularly to obtain mice with deletion of the HO-1 gene in myeloid lineage and are designated as HMOX1WT (intact HO-1) and HMOX1fl/fl (x LysMcre; complete HO-1 deficiency).

### 2.2 CpG-induced hyperinflammation

Secondary hyperinflammatory disease was induced by repeat activation of TLR-9 as previously described [4]. Briefly, 7 to 10-week-old mice were injected with 5 doses of phosphate buffered saline (PBS) or 50 μg of CpG 1826 intraperitoneally every other day for 9 days. On day 10, mice were euthanized 24 hours after the last injection of CpG, blood was collected for complete blood cell counts and serum was isolated for cytokine analysis by enzyme-linked immunosorbent assay (ELISA). Thereafter, mice were euthanized, and the spleens and livers were harvested, with tissue samples formalin-fixed, and then embedded in paraffin for tissue sectioning and slide preparation for histology. Histology was scored by a single blinded pathologist. For IL-10 blockade experiments, 200 ug IgG2b control antibody (clone SFR8) or IL-10 receptor type I (IL-10RI) blocking antibody (referred to as IL-10RB, clone 1B1.3A) (both from Bio X Cell) were injected i.p. on days 0, 2, and 4.

### 2.3 Measuring products of nitrogen and oxygen radicals

Nitrite (as a measure of nitic oxide production) was measured in serum and *in vitro* supernatants using a modified Griess reagent (Sigma-Aldrich, Cat. G4410) as described by Green, et al. [10]. Glutathione GSH/GSSG ratio was measured using Glutathione assay kit (Sigma-Aldrich, Cat. MAK440). Lipid peroxidation was measured using an malondialdehyde (MDA) assay kit (Sigma-Aldrich, Cat. MAK085).

### 2.4 Measuring serum free heme and ferritin

Serum samples were collected from mice treated with CpG and healthy controls. Serum free heme was measured using a heme assay kit per manufacturer instructions (Sigma-Aldrich, Cat. MAK316). Ferritin was measured using a Ferritin ELISA assay from Abcam (cat. ab157713).

### 2.5 Bone marrow derived macrophages and dendritic cells – generation and stimulation

Bone marrow (BM) was isolated from hind limbs of mice and filtered it over a 70-μm strainer to obtain a single-cell suspension, followed by RBC lysis. Bone marrow–derived macrophages (BMDMs) were generated in tissue culture using L929 macrophage colony-stimulating factor–enriched media. Bone marrow-derived dendritic cells (BMDCs) were generated in tissue culture using recombinant GM-CSF (Peprotech, Cat. AF-315-03). Class B CpG 1826 oligonucleotide was synthesized by Integrated DNA Technologies. BMDM/BMDCs were treated with 1 μg/ml of CpG. Incubation times after CpG stimulation are indicated in individual experiments. Recombinant IFN gamma (IFNg) was purchased from Peprotech (Cat. 315-05). Monomethyl Fumarate (MMF) was purchased from Tocris (product no: 4511), resuspended in DMSO. Hemin was purchased from Sigma (Cat. 51280), resuspended in DMSO and incubated with cells at a final concentration of 100 uM.

### 2.6 Cytokine analysis

BD OptEIA ELISA kits were used for analysis of serum from *in vivo* experiments and supernatant from *in vitro* experiments. Product nos. 555256 [IL-12], 555252 [IL-10], 555268 [tumor necrosis factor (TNF)], 555142 [IFNg], and 555240 [IL-6]).

### 2.7 Preparation of oxidized red blood cells

This protocol was adapted from Olonisakin et al. [8]. Briefly, peripheral blood was collected from healthy mice or sick mice treated with CpG where indicated. For *in vitro* studies, RBCs were prepared and used for experiments the same day. Fresh RBCs were collected from healthy C57BL/6 mice.

Stressed/sick RBCs (Sick RBCs) were collected from C57BL/6 mice treated with CpG and washed three times with PBS. Hydrogen peroxide treated RBCs (H2O2 RBCs) were collected from healthy C57BL/6 mice and pretreated with 100 uM hydrogen peroxide. For RBC lysate/ghost preparation: hydrogen peroxide-treated RBCs were centrifuged at 800 g for 10 minutes, washed three times with cold PBS, and lysed with RBC lysis buffer. Lysed RBCs were centrifuged at 30,000 g for 30 minutes at 4°C. RBC ghost pellets were washed three times and reconstituted in PBS. BMDCs were incubated with RBC preparations one hour prior to stimulation with CpG.

### 2.8 Quantitative PCR and mRNA regulation studies

Gene expression in BMDCs was analyzed by quantitative real-time RT-PCR (qPCR) using SYBR Green probes. Briefly, RNA was extracted using Qiagen (Hilden, Germany) RNeasy Mini Kit (Cat. 74106). cDNA was synthesized using SuperScript III First-strand synthesis system (Invitrogen, Cat 18080-051). Pre-synthesized QuantiTect primers from Qiagen were used for il12b. qRT-PCR reaction was measured and processed using StepOne PCR system and software (Applied Biosystems). Relative quantification of gene expression was calculated to housekeeping gene GAPDH. For mRNA studies, cycloheximide (CHX) final concentration of 8 ug/mL was used to arrest protein translation and Actinomycin D (Cell Signaling Technology, Cat. 15021) was used to arrest transcription by treating cells with 10 ug/mL Actinomycin D where indicated. Methods for measuring mRNA turnover was adapted from Lai, et al. [11].

### 2.9 Statistical analysis

Values are shown as the mean ± SEM. Depending on the structure being compared, paired 1-tailed or 2-tailed t-tests or one-way or two-way analyses of variance were performed. Figures were generated and analyzed using GraphPad Prism software, version 7.

## 3. Results

### 3.1 CpG-induced nitrogen radical production is suppressed by monomethyl fumarate in a NRF2 dependent manner

We have previously demonstrated that the small molecule monomethyl fumarate (MMF) is able to ameliorate CpG-induced murine hyperinflammation in a HO1-dependent and HO1-independent manner [4]. Publications from our lab and others have revealed MMF as a potent inducer of transcription factor NRF2, which is well established as a critical sensor and regulator of antioxidant cellular pathways. To test our hypothesis that NRF2 is a regulator of systemic hyperinflammatory states, we first tested whether MMF-induced NRF2 activation could suppress nitrogen radical production by bone marrow-derived macrophages (BMDMs) as measured by a modified Griess reagent. We observed that MMF was able to suppress nitrogen radical production in BMDMs stimulated with CpG alone (Figure 1A) and in BMDMs primed with IFNg prior to CpG stimulation (Figure 1B). Using BMDMs from NRF2 deficient mice demonstrates that MMF-induced suppression of nitrogen radical production occurs in a NRF2-dependent manner.

**Figure 1:**
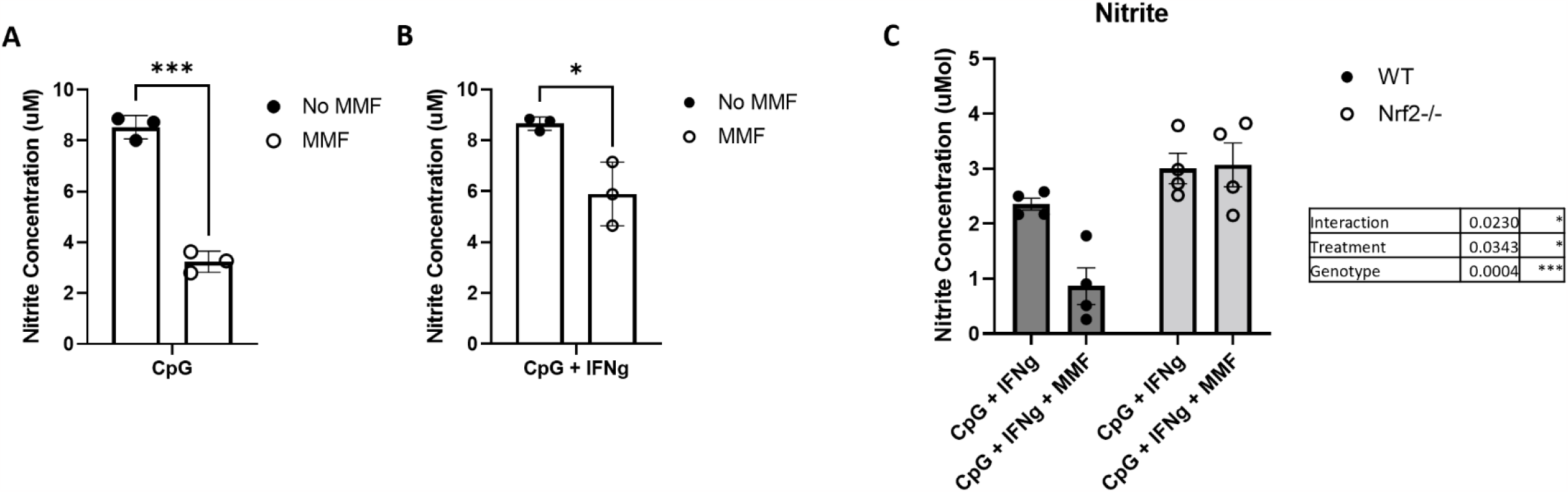
Nitrogen radical production by bone marrow derived macrophages (BMDM) is suppressed by monomethyl fumarate (MMF) in a NRF2 dependent manner. BMDMs were cultured using M-CSF enriched media. **(A, B)** BMDMs were stimulated with CpG with or without IFNg as indicated. Cells were pre-treated with MMF one hour prior to stimulation where indicated. **(C)** Experiments were repeated using BMDMs from wildtype C57BL/6 mice (WT) and from NRF2 -/- mice. Nitrite production was measured by a modified Griess reagent. Figures are representative of three separate experiments. P-values were calculated by pairwise comparison in A and B and two way analysis of variance (ANOVA) in C.

### 3.2 CpG induced hyperinflammation produces increased markers of nitrosative and oxidative distress

NRF2 is a central regulator of redox homeostasis. We therefore first sought to characterize the systemic oxidative environment after induction of CpG-induced hyperinflammation. C57BL/6 (WT) and Nrf deficient (Nrf -/-) mice were injected i.p. with CpG every other day for 10 days (5 doses total) to induce hyperinflammatory disease. We observed elevated concentrations of serum nitrites in CpG treated mice and this was exacerbated by NRF2 deficiency (Figure 2A). We repeated this experiment with HMOX1^*fl/fl*^ LysM Cre mice with the hypothesis that the delta between WT and NRF2 KO nitrite production was secondary to a lack of HO1 activation. We observed that mice lacking HO1 (HMOX1^*fl/fl*^ LysM Cre) overproduced nitrogen radicals to a similar level compared to NRF2 deficient mice (Figure 2B). To further characterize the oxidative environment, we measured the whole blood glutathione levels and lipid peroxidation of RBCs and whole spleen homogenates. As expected, CpG-treated mice had a reduction in the GSH/GSSG ratio (secondary to increased oxidized glutathione) which was worsened by NRF2 deficiency (Figure 2C). Similarly, CpG-treated mice have increased lipid peroxidation in RBC and whole spleen homogenates as measured by MDA assay which is significantly exacerbated in NRF2 -/- mice (Figure 2D).

**Figure 2:**
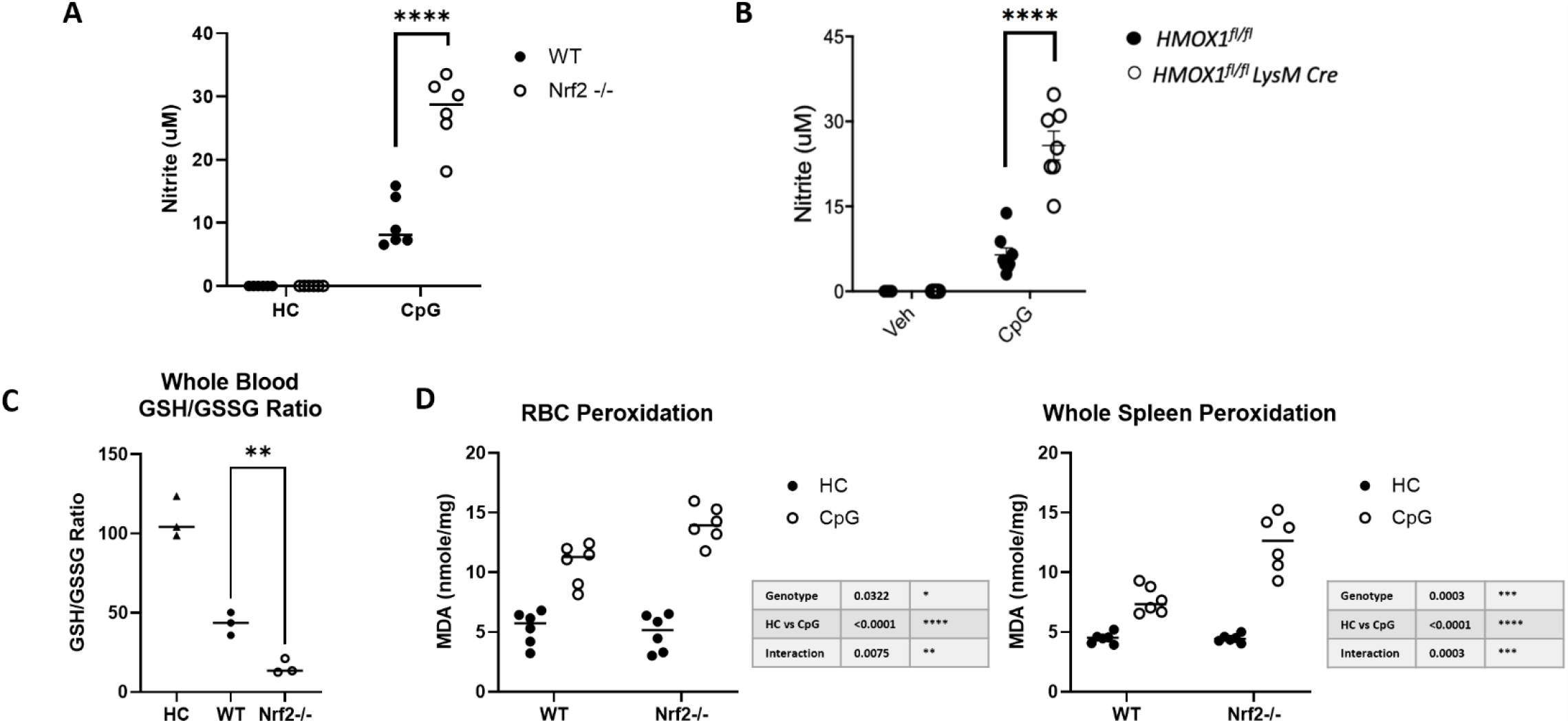
CpG induced hyperinflammation produces elevated markers of nitrosative and oxidative stress *in vivo* which are exacerbated by NRF2 deficiency. **(A, B)** Serum levels of nitrite from healthy controls (HC) and CpG-treated mice were measured by modified Griess reagent. **(C)** Whole blood was collected from healthy controls, wildtype mice treated with CpG, and NRF2 -/- treated with CpG. Levels of reduced glutathione (GSH) and oxidized glutathione (GSSG) were measured and presented as a ratio of GSH/GSSG – with decreased ratios indicative of increased oxidation. **(D)** Lipid peroxidation was measured in whole blood and whole spleen homogenates by MDA assay after treatment with CpG and compared to healthy controls. Figures are representative of two to three separate experiments. P-values were calculated by pairwise comparison in A, B, and C; and by two way ANOVA in D.

### 3.3 Lack of NRF2 exacerbates murine hyperinflammatory disease

Our prior work suggests that targeting NRF2 activation could improve disease markers in murine hyperinflammatory states. We therefore hypothesized that NRF2 deficiency would exacerbate disease in our mouse model of CpG-induced secondary hyperinflammation. C57BL/6 (WT) and Nrf-/- mice were injected i.p. with CpG every other day for 10 days (5 doses total) to induce hyperinflammatory disease. We found that Nrf-/- mice had significant weight stagnation compared to their WT counterparts (Figure 3A). Hepatosplenomegaly, a hallmark of systemic hyperinflammatory disease, is also exacerbated by NRF2 deficiency (Figure 3B). Livers were harvested and histology was scored using a published system [4] by a single blinded pathologist. Scores for hepatic portal inflammation and endothelial activation were significantly higher in the NRF2 deficient mice (Figure 3C). Representative liver histology is shown in Figure 3D. Splenic immune cell populations were analyzed by flow cytometry, revealing no significant difference in immune cell populations in WT versus Nrf-/- mice (supplemental Figure 1).

**Figure 3:**
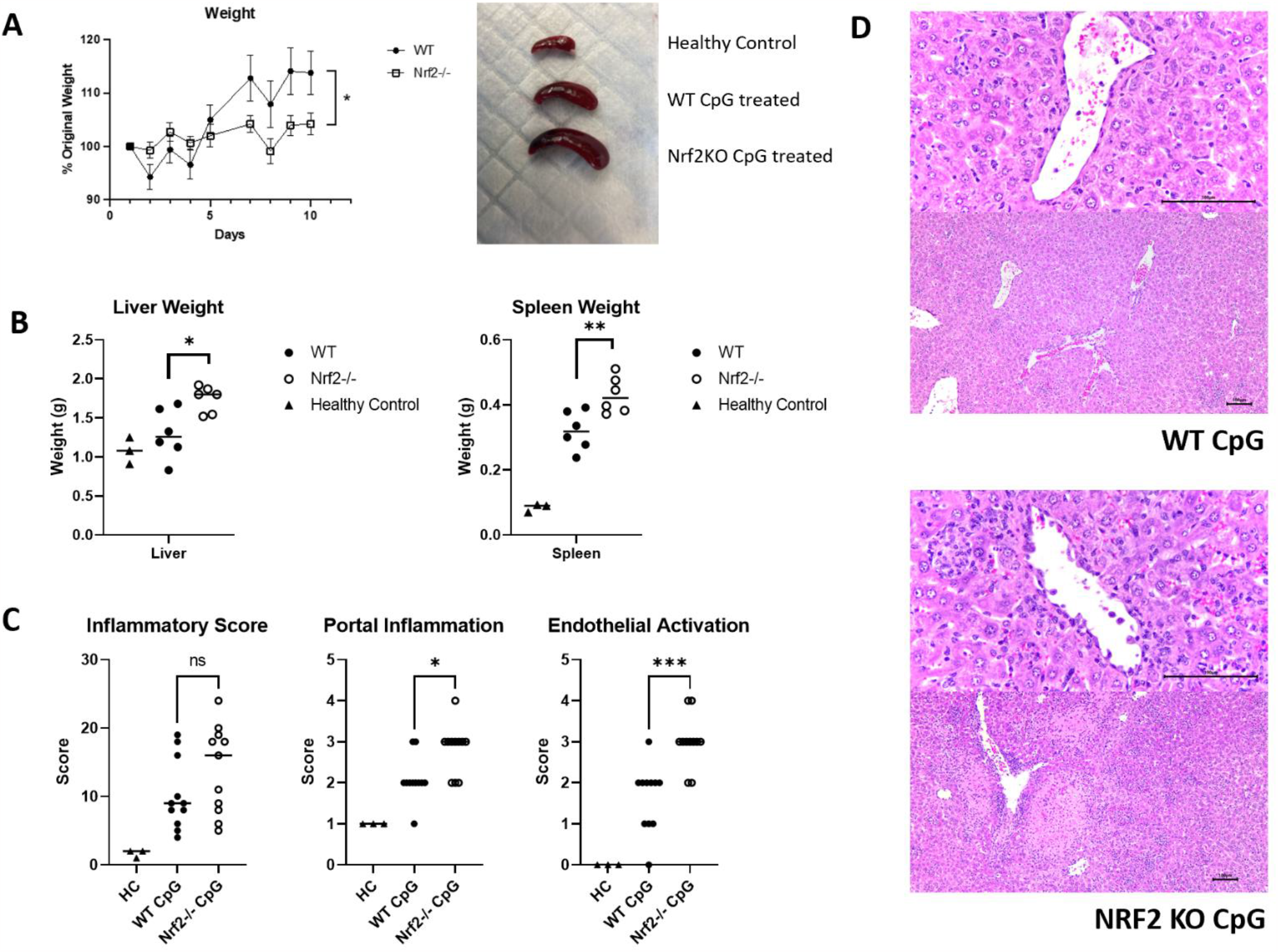
Lack of NRF2 exacerbates end organ damage in CpG induced hyperinflammatory disease. Hyperinflammation was induced in WT and NRF2 -/- mice by repeat i.p. injections of CpG for a total of five doses over a ten day course. **(A)** mouse body weights were tracked daily. **(B)** At the conclusion mice were euthanized. Liver and spleens weights were measured and compared to healthy controls. **(C)** Liver histology was examined and scored by a blind pathologist. Scores for global liver inflammation, portal inflammation, and endothelial activation are shown. **(D)** Representative histologic images are shown. Figures are representative of three separate experiments. P-values were calculated by two way ANOVA in A; and by pairwise comparison in B and C.

We have previously reported that antibody blockade of the IL-10 receptor significantly exacerbates the CpG-induced hyperinflammation toward a more fulminant MAS model [12]. We tested the hypothesis that NRF2 deficiency would further exacerbate IL-10 blockade in this model. Indeed, Nrf-/- mice treated with CpG in the presence of IL-10 receptor blockade have significant weight loss and become more systemically ill compared to their WT counterparts – requiring euthanasia by experimental day three (supplemental Figure 2).

### 3.4 Lack of NRF2 exacerbates cytopenias and increases serum free heme in CpG-induced hyperinflammation

To further characterize the role of NRF2 in hyperinflammatory states we evaluated complete blood counts (CBCs) from WT and NRF2 -/- mice after induction of CpG-induced hyperinflammation. CBCs were analyzed pre-CpG treatment (closed circles, Figure 4A) and post-CpG treatment (open circles, Figure 4A). In both WT and Nrf-/- mice CpG-induced hyperinflammation results in a similar degree of anemia over the course of the experiment and is not significantly different between the two genotypes. In contrast, after CpG-induced hyperinflammation, Nrf-/- mice develop a significantly increased reticulocytosis compared to their WT counterparts – suggesting increased RBC turnover in the Nrf-/- mice. CpG-induced thrombocytopenia was also significantly lower in Nrf-/- mice after CpG-induced hyperinflammation (Figure 4A).

**Figure 4:**
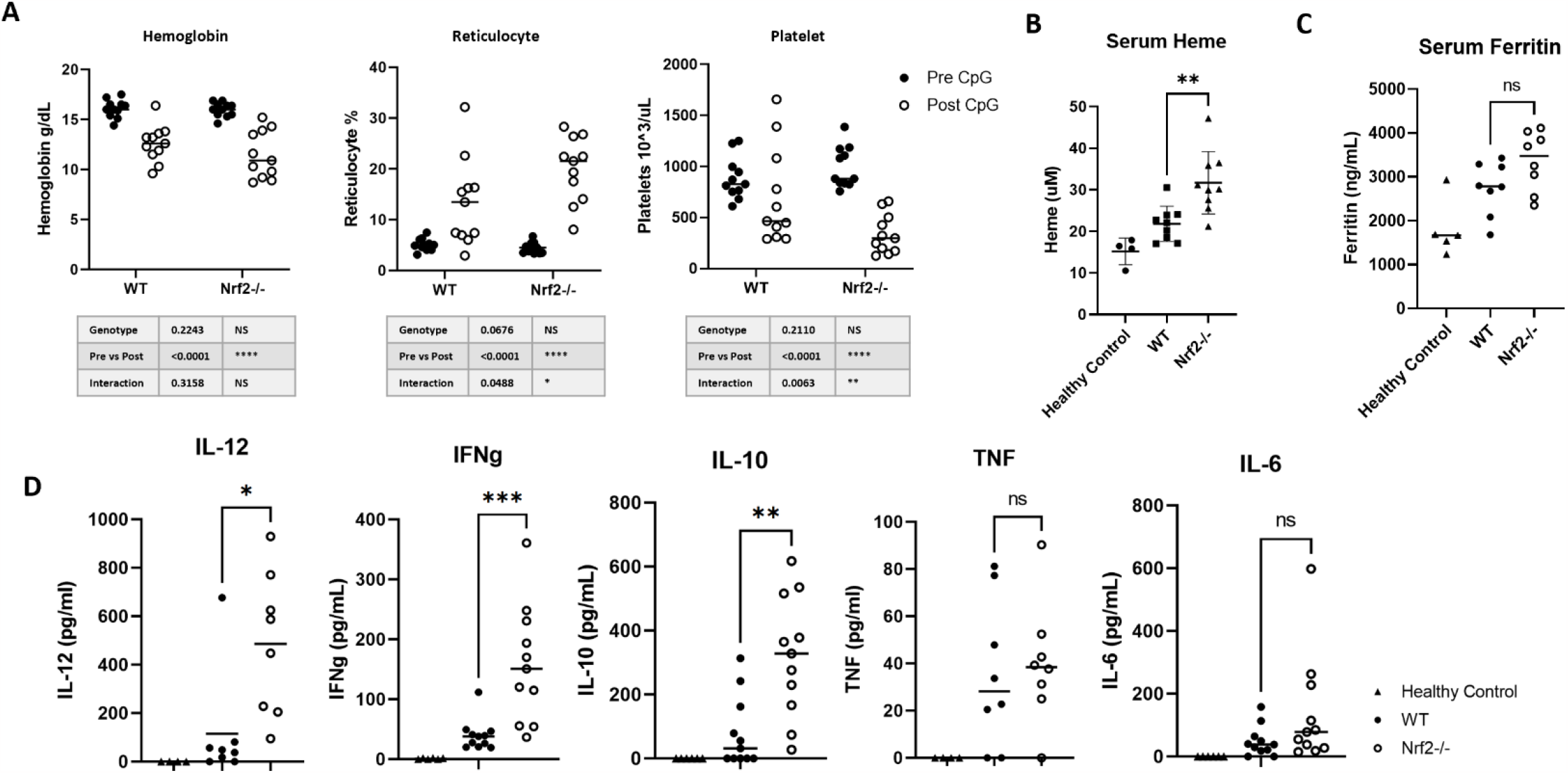
Lack of NRF2 exacerbates cytopenias and hypercytokinemia in CpG induced hyperinflammation. Hyperinflammation was induced in WT and NRF2 -/- mice by repeat i.p. injections of CpG for a total of five doses over a ten day course. Blood samples were collected by cheek bleed. **(A)** Complete blood counts (CBC) and reticulocyte counts were enumerated from WT and NRF2 -/- mice before and after induction of CpG hyperinflammation. **(B)** Serum free heme was measured in WT and NRF2 -/- mice after the induction of CpG hyperinflammation. **(C)** Serum ferritin was measured by ELISA in WT and NRF2 -/- mice after the induction of CpG hyperinflammation. **(D)** Serum cytokines were measured by ELISA in WT and NRF2 -/- mice after the induction of CpG hyperinflammation. P-values were calculated by two way ANOVA in A; and by pairwise comparison in B, C, and D.

Anemia with reticulocytosis is clinically associated with hemolytic disease. We therefore hypothesized that Nrf-/- mice treated with CpG would have elevated levels of serum free heme. Indeed, measurements of serum free heme after CpG-induced hyperinflammation demonstrated elevated levels in Nrf-/- mice (Figure 4B) – suggesting increased RBC damage and hemolysis.

### 3.5 Lack of NRF2 exacerbates IL-12-IFNg-IL-10 hypercytokinemia in CpG-induced hyperinflammation

Hyperferritinemia and hypercytokinemia are clinical hallmarks of hyperinflammatory states like MAS and HLH. We next evaluated serum ferritin and cytokine levels in WT and Nrf-/- mice after CpG-induced hyperinflammation. Both WT and Nrf-/- mice had elevated serum ferritin levels (Figure 4C). Ferritin levels in Nrf-/- mice trended towards increased compared to WT counterparts but was not statistically significant. In contrast, hypercytokinemia was significantly exacerbated in Nrf-/- mice. Cytokines critical to the pathogenesis of MAS/HLH were significantly elevated in Nrf-/- mice after CpG-induced hyperinflammation, namely IL-12, IFNg, and IL-10 (Figure 4D).

### 3.6 Products of oxidized RBCs and Hemin suppress IL-12 production from BMDCs

Hemophagocytosis is another hallmark of hyperinflammatory disease. However, the underlying mechanisms and role of the hemophagocyte in these diseases is still unclear. Work from our lab and others suggests that NRF2 may play a central role in hemophagocyte regulation of systemic hyperinflammation. To explore this hypothesis, we generated BMDCs via *in vitro* cell culture and stimulated them with CpG in the presence of RBCs of varying oxidative states. We measured the IL-12 cytokine response from these cells because prior work has shown that the initial IL-12 response is required for subsequent hyper IFNg state which drives the hyperinflammatory phenotype. Our *in vivo* analysis showed that IL-12, and accordingly IFNg, was elevated in Nrf-/- mice (Figure 4D).

Fresh RBCs were used directly from healthy mice and did not suppress IL-12 protein production by BMDCs stimulated with CpG (Figure 5A). In contrast RBCs from a sick CpG-treated mouse (Sick RBC) and RBCs treated with hydrogen peroxide (H2O2 RBC) were both able to suppress IL-12 production in a NRF2 dependent manner (Figure 5A). This observation suggests that erythrophagocytosis of oxidized RBCs suppresses proinflammatory cytokine production through an NRF2-dependent mechanism.

**Figure 5:**
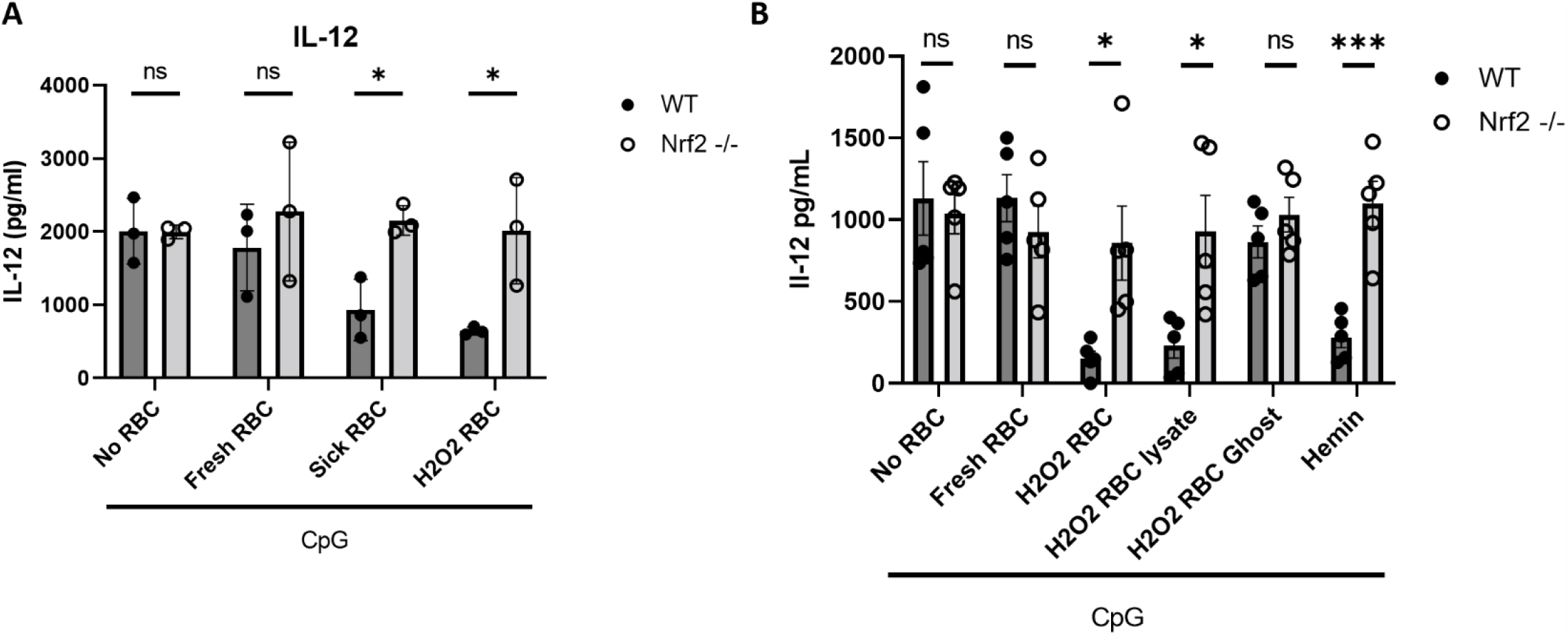
Products of oxidized RBCs and Hemin suppress IL-12 production from bone marrow derived DCs *in vitro*. Bone marrow derived dendritic cells (BMDCs) were cultured using rGM-CSF. **(A)** BMDCs from WT and NRF2-/- mice were incubated in the absence or presence of red blood cells (RBCs) followed by stimulation with TLR-9 agonist CpG. Fresh RBCs were collected by cheek bleed and used the same day. Sick RBCs were collected from a sick CpG-treated hyperinflammatory mouse. H2O2 RBCs are fresh RBCs treated with 100 uM hydrogen peroxide. IL-12 production was measured by ELISA. **(B)** BMDCs from WT and NRF2-/- mice were incubated in the absence or presence of RBCs and RBC products followed by stimulation with CpG. H2O2 RBCs were separated into intracellular lysate and plasma membrane ghost where indicated. Hemin was resuspended in DMSO and incubated with cells at a final concentration of 100 uM. Equivalent DMSO was added to appropriate controls. IL-12 production was measured by ELISA. P-values were calculated by pairwise comparison.

NRF2 and its binding partner Keap1 sense cellular redox distress. We next asked if NRF2 activation by oxidized RBCs was mediated by detection of the oxidized red cell lipid membrane or by cellular contents, namely heme. To test this, we lysed hydrogen peroxide-treated RBCs and separated the cellular contents (lysate) from the red cell plasma membrane (ghost). We then stimulated BMDCs with CpG after incubation with lysate or ghost. Similar to whole oxidized RBC preparations, the cellular lysate was able to suppress IL-12 in a NRF2-dependent manner, but the ghost was not able to suppress IL-12 production (Figure 5B). We hypothesized that NRF2 activation by RBC lysates was mediated by intracellular heme. To test this hypothesis we used hemin, an endogenously produced oxidized product of heme. We observed that hemin was also able to suppress IL-12 production in a NRF2-dependent manner (Figure 5B).

### 3.7 NRF2 suppression of IL-12 mRNA transcription is independent of protein translation and mRNA degradation

IL-12 plays a critical role in the initiation and exacerbation of the positive feedback look with IFNg which propagates hyperinflammatory states. How erythrophagocytosis and heme-mediated activation of NRF2 suppresses IL-12 production is thus of critical clinical importance. IL-12 is a heterodimer encoded by two genes, *il12a* and *il12b*. Further analysis of BMDCs stimulated with CpG in the presence of hemin demonstrates hemin is able to significantly decrease *il12b* mRNA levels in a NRF2 dependent manner, indicating that NRF2 suppresses IL-12 mRNA at the transcriptional level (Figure 6A). However, *il12b* does not have overt NRF2 binding sites in its regulatory regions (Supplemental Table 1), as analyzed by MotifMap [13, 14]. Together this suggests that NRF2 either has a non-canonical binding motif on the *il12b* locus or that NRF2 has an indirect effect on IL-12 regulation.

**Figure 6:**
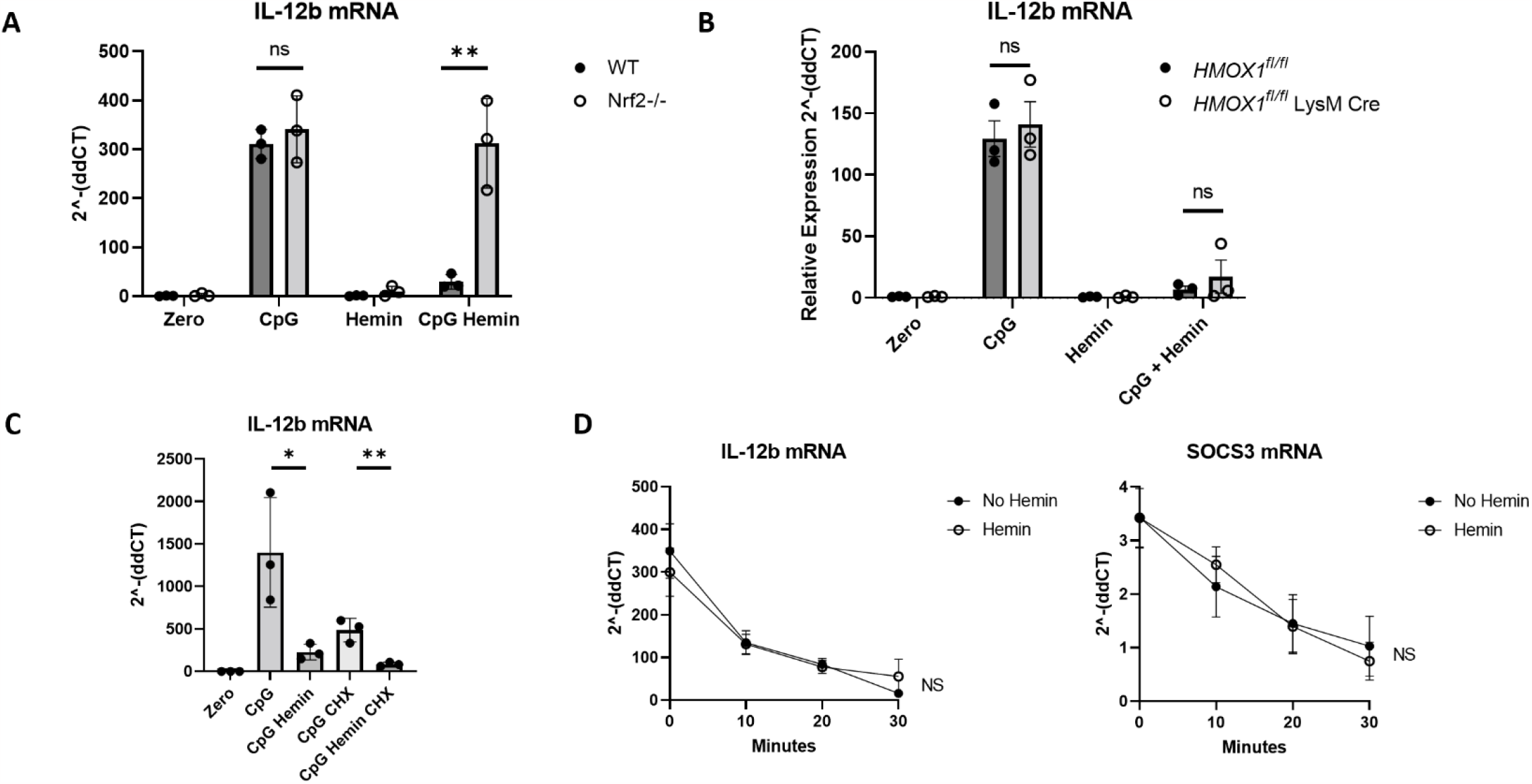
NRF2 suppression of IL-12 mRNA transcription is independent of protein translation and mRNA degradation. Bone marrow derived dendritic cells (BMDCs) were cultured using rGM-CSF. **(A)** BMDCs from WT and NRF2 -/- mice were treated with CpG and/or hemin where indicated. Hemin was added one hour prior to CpG stimulation. IL-12b mRNA transcripts were measured by qRT-PCR. **(B)** BMDCs from HMOX1WT (intact HO-1) and HMOX1fl/fl (x LysMcre; complete HO-1 deficiency) were stimulated with CpG and/or hemin where indicated. IL-12b mRNA transcripts were measured by qRT-PCR. **(C)** BDMCs from WT mice were pre-treated with cycloheximide (CHX) final concentration of 8 ug/mL followed by stimulation with CpG and/or hemin. IL-12b mRNA transcripts were measured by qRT-PCR. **(D)** BDMCs from WT mice were pre-treated with actinomycin D 10 ug/mL and stimulated with CpG with or without hemin. Cells treated with hemin were compared to those not treated with hemin. IL-12b and SOCS3 (control) transcripts were measured by qRT-PCR. P-values were calculated by pairwise comparison in A, B, and C; and by two way ANOVA in D.

We first tested if NRF2 target HO-1 might have an influence on IL-12 mRNA expression, as we have found that HO-1 has a direct effect on IL-10 production. We observed that BMDCs lacking HO1 (HMOX1^*fl/fl*^ LysM Cre) were able to suppress IL-12 mRNA, suggesting that IL-12 suppression is HO-1 independent (Figure 6B).

We next hypothesized that NRF2 suppression of IL-12 requires NRF2 to transcribe a new product and subsequently suppressing IL-12. To test if IL-12 mRNA expression is dependent on new protein translation we first pretreated BMDCs with cycloheximide to block protein translation. We observed that hemin was still able to suppress *il12b* levels in the absence of new protein production (Figure 6C). This is supported by our observations that hemin can suppress IL-12 production when added to culture at the same time as CpG activation, allowing no time for protein translation to occur prior to stimulation.

We next tested if hemin treatment increases *il12b* mRNA instability and therefore increases mRNA degradation. Actinomycin D was used to arrest mRNA transcription. *il12b* mRNA was measured at 0, 10, 20, and 30 minutes after hemin stimulation. SOCS3 was used as a positive control for mRNA turnover [11]. We found no difference in *il12b* mRNA degradation when treated with hemin (Figure 6D). Together these data demonstrate hemin is a potent suppressor of IL-12 protein production and IL-12 mRNA by innate immune cells, most likely acting as a transcriptional repressor through a non-ARE binding mechanism.

## 4. Discussion

As hypothesized by Metchnikoff with their initial discovery in 1882, mononuclear phagocytes have fundamental roles in both innate immunity and tissue homeostasis. Their innate immune function is in part mediated by the respiratory burst – the rapid production of oxygen radicals to kill microbes. But the oxidative damage that subsequently occurs is then regulated and processed by the same mononuclear phagocytic system. These hemostatic functions of mononuclear cells include clearance of apoptotic cell debris, RBC recycling, metabolic regulation, and tissue homeostatic surveillance [15]. Inflammatory and hemostatic processes therefore intersect at a central point – redox homeostasis and iron metabolism.

Hyperinflammatory diseases, like MAS and HLH, are maladaptive processes where normal inflammatory mechanisms become unregulated. It stands to reason that an acute hyperinflammatory state would be one in which the oxidative environment is dysregulated. The presence of hemophagocytosis and other markers of iron dysregulation in MAS/HLH are important clues to the underlying pathophysiology. Despite their recognition for multiple decades, the role of the hemophagocyte in the pathogenesis and regulation of hyperinflammatory syndromes is not well understood. In this report we begin to explore the critical intersection between red blood cell biology, hemophagocytosis, and immune regulation in hyperinflammatory conditions.

We demonstrate that murine hyperinflammation is associated with systemic nitrosative and oxidative distress as measured by serum nitrites and cellular lipid peroxidation which is exacerbated by lack of NRF2. We found that NRF2 deficient mice have worse clinical evidence of RBC damage and lysis after induction of CpG hyperinflammation. In addition to increased RBC peroxidation, NRF2 -/- mice develop a significantly elevated reticuolocytosis in response to their anemia after CpG-induced hyperinflammation. This suggests the hematopoietic compartment is working harder in NRF2 deficient mice to maintain their hemoglobin. RBC lipid peroxidation, elevated reticulocytosis, and elevated serum free heme all suggest RBC damage and increased RBC turnover – which we show is exacerbated by NRF2 deficiency. The reason for increase RBC damage is likely multifactorial. NRF2 deficiency results in a more oxidative environment, but RBCs from NRF2 -/- mice also have a lowered resistance to oxidative distress as demonstrated by Lee et al. [16].

Using the CpG-induced mouse model of hyperinflammation we found that NRF2 deficient mice have significantly worse organomegaly and liver pathology – suggesting an important regulatory role for NRF2 in hyperinflammatory disease. This observation is consistent with our prior work showing that NRF2 activator MMF can ameliorate some aspects of hyperinflammatory disease in this model. We found that NRF2 -/- mice have significantly worse hypercytokinemia – notably favoring cytokines canonical to hyperinflammatory states, namely IL-12, IFNg, and IL-10. Because NRF2 seems to specifically regulate this pathologic cytokine axis, it becomes an attractive therapeutic target for clinical cytokine suppression.

To further explore the relationship between RBC damage, hemophagocytosis, and cytokine suppression we utilized BMDCs *in vitro* to interrogate IL-12 production and mRNA processing - the first key cytokine in hyperinflammatory positive feedback loops. We demonstrate that washed RBCs from a sick CpG-treated mouse can suppress proinflammatory IL-12 production and that this observation is phenocopied using RBC treated with hydrogen peroxide. We separated hydrogen peroxide-treated RBCs into lysates and plasma membrane ghosts – finding that only lysates were able to suppress IL-12 production. Subsequently we demonstrate that hemin (ferric chloride heme, an oxidized form of heme) is sufficient for IL-12 suppression.

These observations are consistent with prior literature demonstrating RBC products and heme suppress proinflammatory cytokine production in a NRF2-dependent manner [8, 9]. However, the IL12b locus does not have overt NRF2 binding sites (Supplemental Table 1). We therefore investigated possible indirect ways NRF2 might regulate IL-12 mRNA expression. We show that hemin’s ability to suppress IL-12 production is independent of protein translation and mRNA turnover – ruling out a translated intermediate and accelerated mRNA degradation, respectively. Although it has previously been shown that NRF2 directly suppresses some cytokines like IL-6 [17], it remains unclear how NRF2 is suppressing IL-12. We hypothesize that NRF2 has binding partners and/or binding sites that remain yet undiscovered.

Together our work demonstrates that transcription factor NRF2 plays a central role in hyperinflammatory disease as a critical regulator of proinflammatory cytokine production. We suggest that RBC damage and ingestion by myeloid cells contributes to a negative feedback loop in which heme activates NRF2, subsequently suppressing proinflammatory cytokine production. Future work is planned to build on the observations reported here. These include elucidating the exact mechanisms by which NRF2 suppresses IL-12 production, confirming the role of NRF2 in a primary model of perforin deficient HLH, and investigating therapeutic strategies targeting NRF2 in human hyperinflammatory disease.

## Acknowledgements

This work was supported by Nancy Taylor Foundation for Chronic Diseases. The graphic abstract was created with BioRender.

## Authorship Contributions

PG: Experimental design, investigation, data acquisition and analysis, manuscript drafting; EE, GF, CB, JK, JW, CJ, NC, ML, TD: Experimental processing, assay processing, animal assistance; PK: Histopathology soring; ML: Scientific mentoring; EB: Experimental design, data interpretation, manuscript writing and revision.

