## Supplemental Figures for "Transcription Factor NRF2 is Activated by Erythrophagocytosis of Oxidized Red Blood Cell Products and Suppresses the IL-12-IFNg-IL-10 Axis in a Murine Model of Hyperinflammatory Disease"

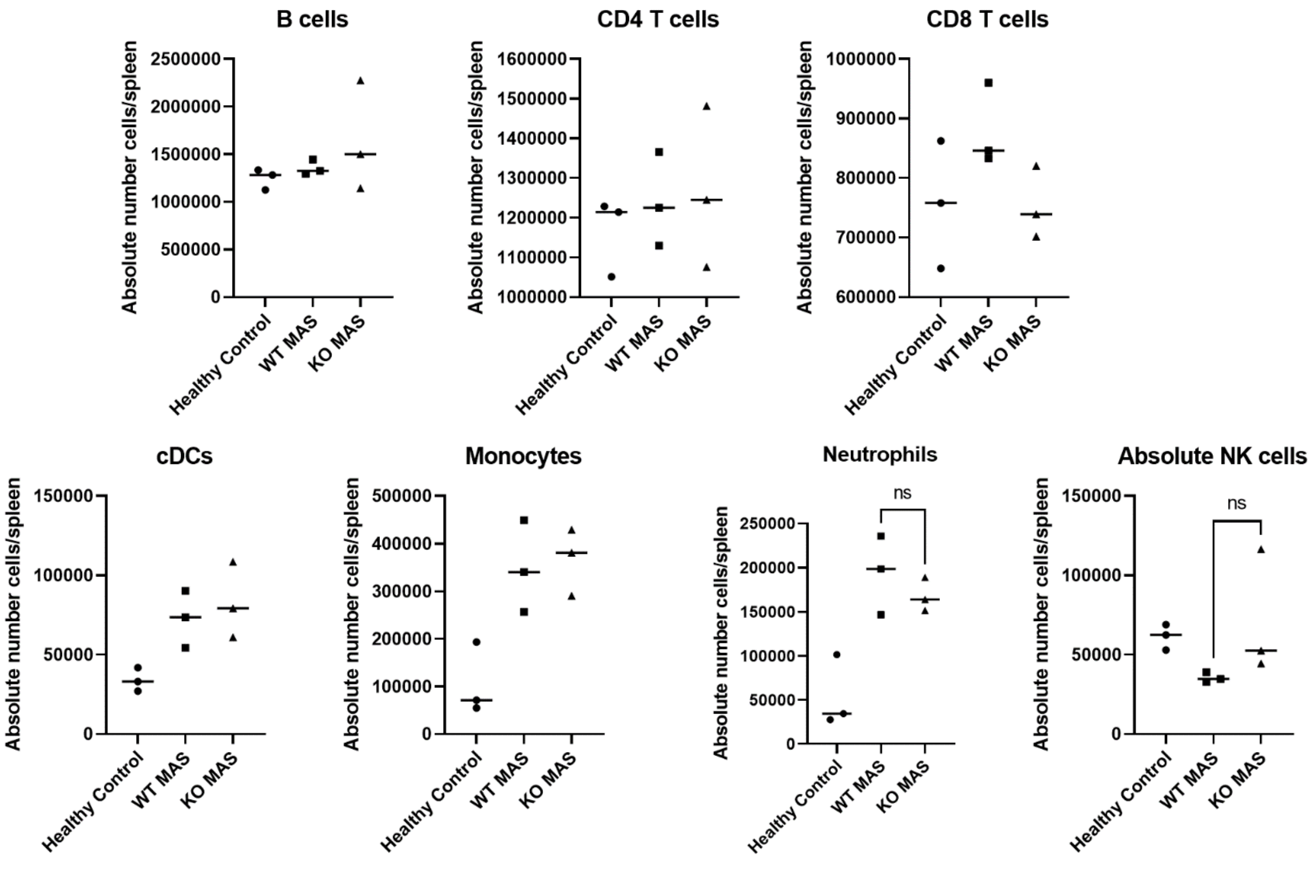


**Supplemental Figure 1: Flow cytometric analysis of immune cell populations in the spleen of mice with CpG induced hyperinflammation.** Hyperinflammation (MAS) was induced in WT and NRF2 -/- mice by repeat i.p. injections of CpG for a total of five doses over a ten day course. Spleens were harvested after CpG-induced hyperinflammation and processed for flow cytometry. Immune cell populations were enumerated by flow cytometry.

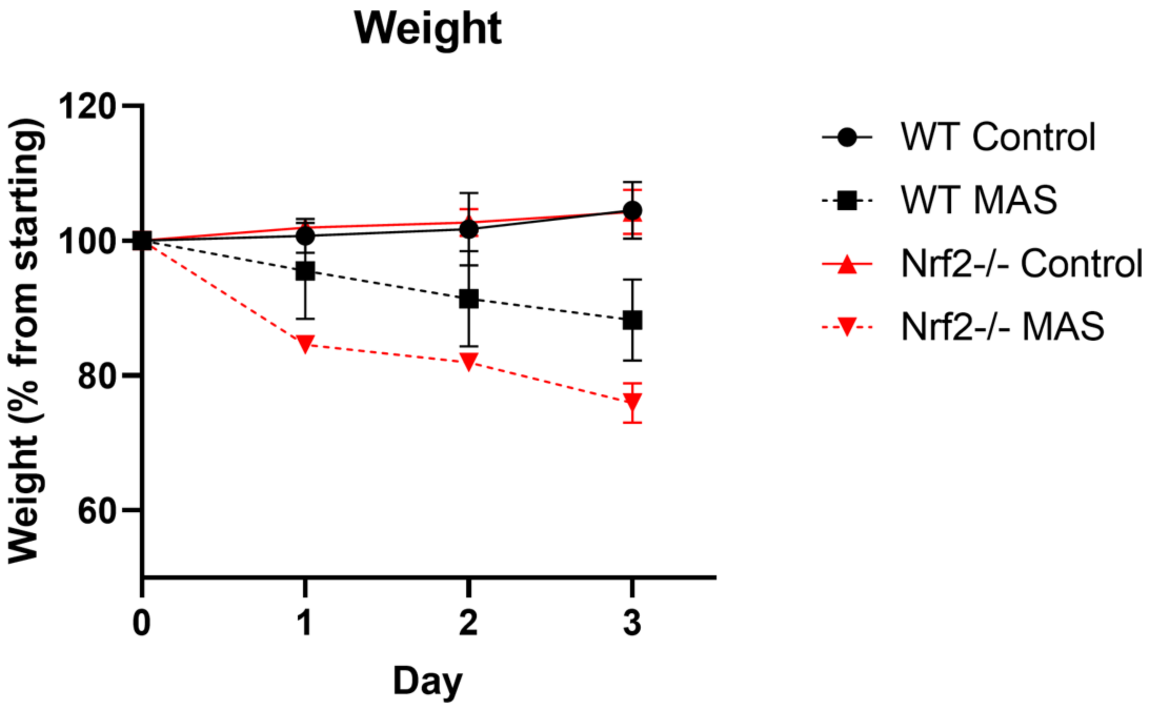


**Supplemental Figure 2: Daily weights of mice treated with CpG and anti-IL-10R.** Hyperinflammation (MAS) was induced in WT and NRF2 -/- mice by repeat i.p. injections of CpG in the presence or absence of blocking antibody to the IL-10 receptor. Mouse body weight was measured daily. The experiment was terminated for significant weight loss in the CpG-treated NRF2 -/- group.

| NQO1 | | HMOX1 | | IL12b | |
|---|---|---|---|---|---|
| sequence | TF-name | sequence | TF-name | sequence | TF-name |
| GTGACTCAG | AP-1 | AGGTCAC | FXR | AGGTCAC | FXR |
| GTGACTCAG | AP-1 | TGCTGAGTCGC | NF-E2 | GCTGAC | MAFB |
| AGTGACTCAGC | AP-1 | CAGCTG | Neuro D | TCAGCAG | MAFA |
| AGTGACTCAGC | AP-1 | CAGCTG | Neuro D | AGTTTCA | IRF8 |
| AGTGACTCAGC | AP-1 | ATGACGCAGCA | **NFE2L2** | GGGAAGTTCC | NF-kappaB |
| TGCTGAGTCAC | NF-E2 | CTGCTGCGTCATG | Nrf-2 | GGGGGAAGTTCCTG | NF-kappaB |
| GTGACTCAGCA | **NFE2L2** | CAAACATGACGCAGCAGAAATG | TCF11:MafG | GGTGGGGGAAGTTCCT | NF-kappaB |
| CAGTGACTCAGCA | AP-1 | TTTGGGAGG | Lyf-1 | CCGCCGACAGAGGTCGCTG | CTCF |
| TCAGCAG | MAFA | GTGACTCAG | AP-1 | GCCGCCGACAGAGGTCGCTG | CTCF |
| CTGCTGAGTCACT | Nrf-2 | GTGACTCAG | AP-1 | CCTCTGTCGGCGGCTCTGG | NRSF |
| TCACAGTGACTCAGCAGAATCT | TCF11:MafG | TGCTGAGTCAC | NF-E2 | CCCAGAGCCGCCGACAGAGGT | NRSF |
| TGCGCAC | MTF1 | GTGACTCAGCA | NFE2L2 | TTCATTTGCACAC | OCT-x |
| CTTCCTG | ETS2 | TTGCTGAGTCACC | Nrf-2 | GGGGAATTTCAA | NF-kappaB |
| | | GGTGACTCAGC | AP-1 | GGGGAATTTC | RelB:p52 (NF-kappaB) |
| | | GGTGACTCAGC | AP-1 | CAGCAACAGCAG | Myf |
| | | GGTGACTCAGC | AP-1 | TCCAGTACCAGCAACAGCAGC | REST |
| | | GGTGACTCAGC | AP-1 | TCCAGTACCAGCAACAGCAGC | NRSE |
| | | GCTGAGTCACC | Bach2 | TCCAGTACCAGCAACAGCAGC | NRSF |
| | | TGGTGACTCAGCA | AP-1 | CAGCTG | Neuro D |
| | | CACTGGTGACTCAGCAAAATCT | TCF11:MafG | CAGCTG | Neuro D |
| | | AGTTTCA | IRF8 | CAGCTG | Neuro D |
| | | AAGTTTCACTTCCTC | ISGF-3 | CAGCTG | Neuro D |
| | | TTCTAGAAGTTTC | HSF | TGCTTCCAGGAATCC | STAT1 |
| | | GAAACTTCTAGAA | HSF2 | TGCTTCCAGGAATCC | STAT5A (homodimer) |
| | | TTTCTAGAAGTTTCACT | HSF1 | GGATTCCTGGAAGCA | STAT5A (homodimer) |
| | | ATGCTGATTCAGC | Nrf-2 | GGATTCCTGGAAGCA | STAT1 |
| | | GCTGAC | MAFB | GATGACTCACC | AP-1 |
| | | TCAGCAG | MAFA | TTTCCAC | NFAT2 |
| | | CAGCTG | Neuro D | CAGCTG | Neuro D |
| | | CAGCTG | Neuro D | CAGCTG | Neuro D |
| | | CACGTGG | USF1 | | |
| | | CACGTGG | Myc | | |
| | | CACGTGA | USF1 | | |
| | | TCACGTGG | CLOCK:BMAL | | |
| | | CCACGTGA | CLOCK:BMAL | | |
| | | CCACGTGA | USF | | |
| | | GCCCACGTGA | MAX | | |
| | | CCCACGTGAC | USF | | |
| | | GTCACGTGGG | USF | | |
| | | GCCCACGTGACC | N-Myc | | |
| | | CCACGTGACC | Ebox | | |
| | | GGTCACGTGG | MAX | | |
| | | GCCCACGTGACC | USF | | |
| | | GGCCCACGTGACCC | MAX | | |
| | | GGGTCACGTGGGCC | MAX | | |
| | | GGCCCACGTGACCC | USF | | |
| | | GGGTCACGTGGGCC | USF | | |
| | | CGGGTCACGTGGGCCA | Arnt | | |
| | | CCTGGCCCACGTGACCCGC | ER | | |
| | | GGCGGGTCACGTGGGCCAGG | c-Myc:Max | | |
| | | CCTGGCCCACGTGACCCGCC | Arnt | | |
| | | GGCGGGTCACGTGGGCCAGG | Arnt | | |
| | | CCTGGCCCACGTGACCCGCC | c-Myc:Max | | |
| | | TCAGCAG | MAFA | | |
| | | CAGCTG | Neuro D | | |
| | | CAGCTG | Neuro D | | |

**Supplemental Table 1: Predicted transcription factor binding sites for NQO1, HMOX1, and IL-12b.** MotifMap was used to probe for NRF2 binding sites in the IL-12b gene, with NQO1 and HMOX1 as controls. NQO1 and HMOX1 revealed NRF2 binding sites, but IL-12b revealed no putative binding sites.
